# Giant magnetoresistive biosensors for real-time quantitative detection of protease activity

**DOI:** 10.1101/2020.01.26.917476

**Authors:** Sandeep Adem, Sonal Jain, Michael Sveiven, Xiahan Zhou, Anthony J. O’Donoghue, Drew A. Hall

## Abstract

Proteases are enzymes that cleave proteins and are crucial to physiological processes such as digestion, blood clotting, and wound healing. Unregulated protease activity is a biomarker of several human diseases. Synthetic peptides that are selectively hydrolyzed by a protease of interest can be used as reporter substrates of unregulated protease activity. We developed an activity-based protease sensor by immobilizing magnetic nanoparticles (MNP) to the surface of a giant magnetoresistive spin-valve (GMR SV) sensor using peptides. Cleavage of these peptides by a protease, releases the magnetic nanoparticles resulting in a time-dependent change in the local magnetic field. Using this approach, we detected a significant release of MNPs after 3.5 minutes incubation using just 4 nM of the cysteine protease, papain. In addition, we show that proteases in healthy human urine do not release the MNPs, however addition of 20 nM of papain to the urine samples resulted in a time-dependent change in magnetoresistance. This study lays the foundation for using GMR SV sensors as a platform for real-time quantitative detection of protease activity in biological fluids.

---

Proteases play an important role in various physiological activities such as food digestion^1^, wound healing^2^, immune function^3^, and intracellular protein turnover^4^. These enzymes cleave between amino acids in proteins and peptides and are the largest class of post-translational modifying enzymes in the human proteome^5^. In cancer and neurodegeneration, unregulated proteolysis can occur when excess proteases are present at the site of disease or alternatively when the endogenous inhibitors are lacking^6,7^. Detection and quantitation of proteases in biofluids can provide a greater understanding of diagnosis and staging of these diseases^8,9^. For example, increased levels of the prostate-specific antigen (PSA) protease in blood is correlated with prostate cancer. This protease is quantified by an enzyme-linked-immunosorbent assay (ELISA)^10^. However, since proteases are catalytically active, there is considerable interest in quantifying enzyme activity rather than protein levels^11,12^. Recently, Ivry and colleagues showed that activity from two aspartic acid proteases, gastricin and cathepsin E, is significantly increased in pre-malignant pancreatic cyst fluid when compared to benign cyst fluid^13^. Development of an assay to screen patient-derived cyst fluid for these protease activities has the potential to stratify patients for surgical intervention or surveillance. In addition, high levels of protease activity occurs in sputum of patients with chronic obstructive pulmonary disease^14,15^, cystic fibrosis^16,17,18^, and in non-healing wounds^19,20^. Rapid and accurate quantification of proteases in biofluids such as plasma, urine, sputum, saliva and wound fluid will allow clinicians to make important decisions about the treatment regimen.

To develop an activity-based protease biomarker of disease, it is essential to utilize a substrate that is efficiently cleaved by the protease of interest. Protease assays can be categorized as either homogenous or heterogeneous assays^21^. In a homogenous assay, the substrate and enzyme are both present in solution and generally utilize fluorescent or colorimetric peptide substrates that are illuminated upon protease cleavage^22,23^. In a heterogeneous assay, the enzyme is in solution while the substrate is immobilized. Heterogeneous protease assays have utilized electrochemical detection^24,25^, surface plasmon resonance (SPR)^26^, surface-enhanced Raman spectroscopy (SERS)^27^, ELISA, and liquid crystal technology^28^ and these assays have been developed for detection of enzyme activity for trypsin, caspase-3, and matrix metallopeptidases^21^.

In this study, we developed a heterogeneous protease assay that uses giant magnetoresistive spin-valve (GMR SV) sensors. These sensors transduce changes in the local magnetic field into electrical signals and have been used as the read head in hard disk drives^29,30^, current sensors^31,32^, magnetic memory^33^ and biosensors^34,35^. Their operation is rooted in quantum mechanics, exhibiting a phenomenon known as spin-dependent scattering^36^, wherein the device’s resistance is proportional to the magnetic field. These sensors are highly scalable, can be produced at low cost, and manufactured in high volume. There are several advantages to using magnetic-based sensors for a protease assay over homogenous and other heterogeneous assay formats. First, clinical samples do not contain any magnetic content. Therefore, these samples have intrinsically low background signal, enabling high sensitivity. In comparison, optical assays face problems such as autofluorescence^37^ and label-bleaching^38^ that produce an undesired background signal. Second, the GMR sensors can be arrayed for multiplexed detection in a single assay without the need for optical scanning. Third, magnetic sensors are insensitive to the sample matrix, allowing them to be used with a wide variety of samples with minimal sample preparation^39^, making then convenient for use in point-of-care (POC) or point-of-use (POU) settings^40^. Lastly, the sensors continuously quantify the local magnetic field changes enabling real-time monitoring of the assay.

In previous applications of GMR SV biosensor technology, recruitment of magnetic nanoparticles (MNPs) to the surface via an antibody-antigen interaction was quantified in real-time^41^. However, in this study, we use this technology in reverse and without the use of antigens or antibodies. MNPs are first tethered to the magnetic sensor surface using a peptide, therefore creating an environment with high magnetoresistance (MR) signal. The peptide sequence is designed to be a substrate for the cysteine protease, papain. Upon addition of this enzyme, the peptide sequence is hydrolyzed, causing the MNPs to be released from the proximity of the sensor (Fig. 1). This results in a decrease in MR that can be monitored in real-time. When a fixed amount of peptide-MNP substrate is present, the release rate correlates with the protease concentration.

**Figure 1.**
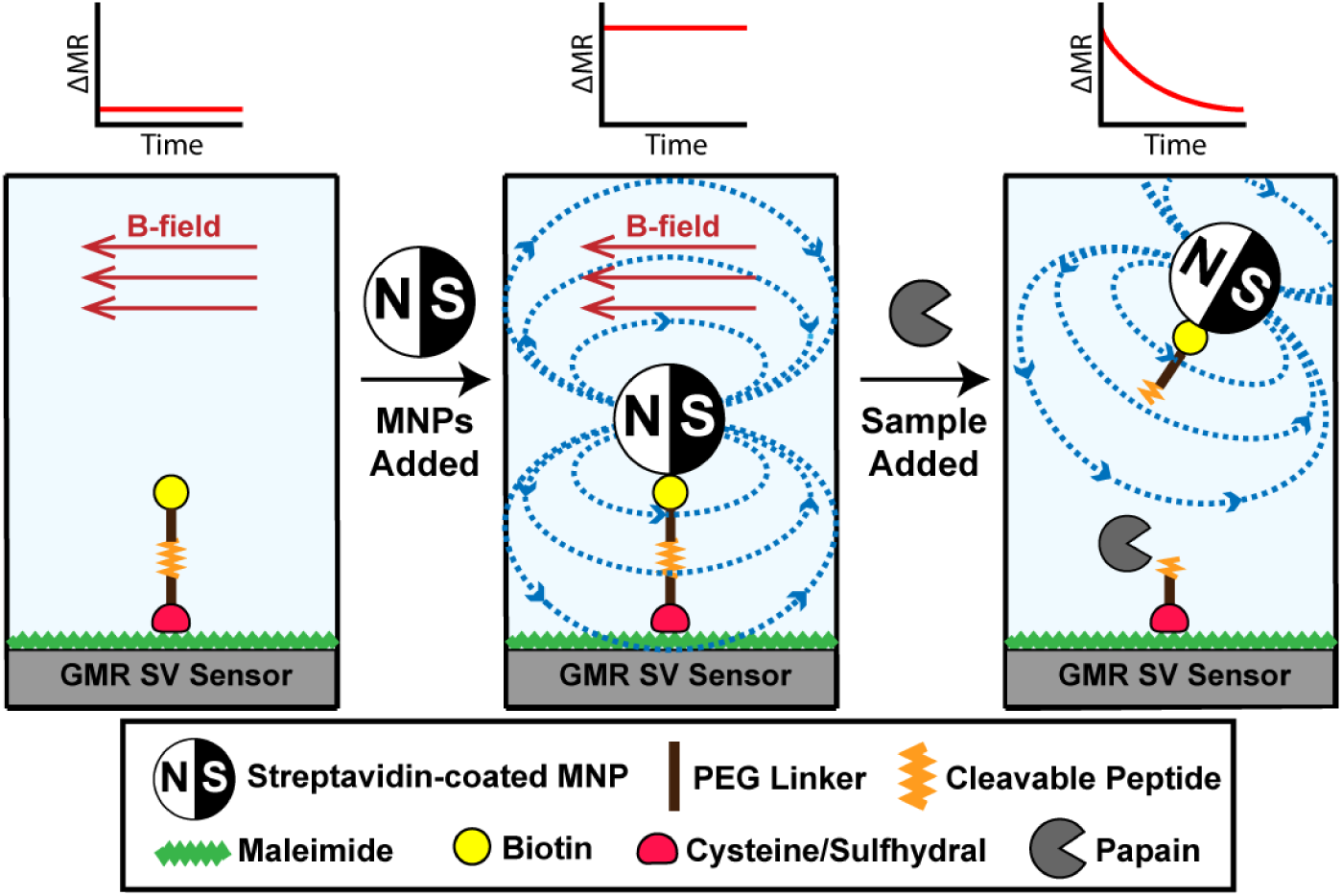
Illustration of the magnetic detection scheme for protease activity. A biotinylated peptide is immobilized on the GMR SV sensors and placed in a magnetic field. Addition of streptavidin coated MNPs causes an increase in magnetoresistance (MR) as they are orientated close to the sensor surface via the streptavidin-biotin interaction. When a biofluid sample containing a protease is added, cleavage of the peptide causes a time-dependent change in the MR as the MNPs are enzymatically released from the sensor surface.

## RESULTS

### Design of papain peptide substrate for heterogeneous assay

Papain is a well characterized cysteine protease related to several disease-associated proteases such as cathepsin K^42^ and cruzain^43^. This enzyme is optimally active between pH 5.5 and 7.5 and irreversibly inactivated by the epoxide inhibitor, E-64^44^. We used this enzyme as a model system to test and validate our proof-of-concept heterogeneous GMR SV based protease assay. To identify a peptide sequence that is efficiently cleaved by papain, the enzyme was previously combined with an equimolar mixture of synthetic tetradecapeptides and cleavage products were quantified by mass spectrometry^45^. Using these data, we identified a tetradecapeptide substrate, KWLIHPTFSYnRWP that was rapidly cleaved between S-Y and Y-n, where lowercase ‘n’ corresponds to the non-natural amino acid, norleucine. We synthesized the C-terminal region of this peptide that contained the sequence TFSYnRWP, corresponding to the amino acids that surround the cleavage sites. However, to make this sequence compatible with the heterogeneous assay format, Biotin-PEG_36_ and PEG_12_-Cys were included on the *N*-terminus and C-terminus, respectively (Fig. 2a). The function of the C-terminal cysteine residue is to facilitate immobilization of the Biotin-PEG_36_-TFSYnRWP-PEG_12_-Cys molecule (hereafter referred to as “peptide”) to a maleimide coated surface through a covalent thioether linkage. The function of the *N*-terminal biotin group is to bind a streptavidin coated reporter element such as streptavidin-marina blue (SA-MB) or streptavidin-coated MNP (SA-MNP).

**Figure 2.**
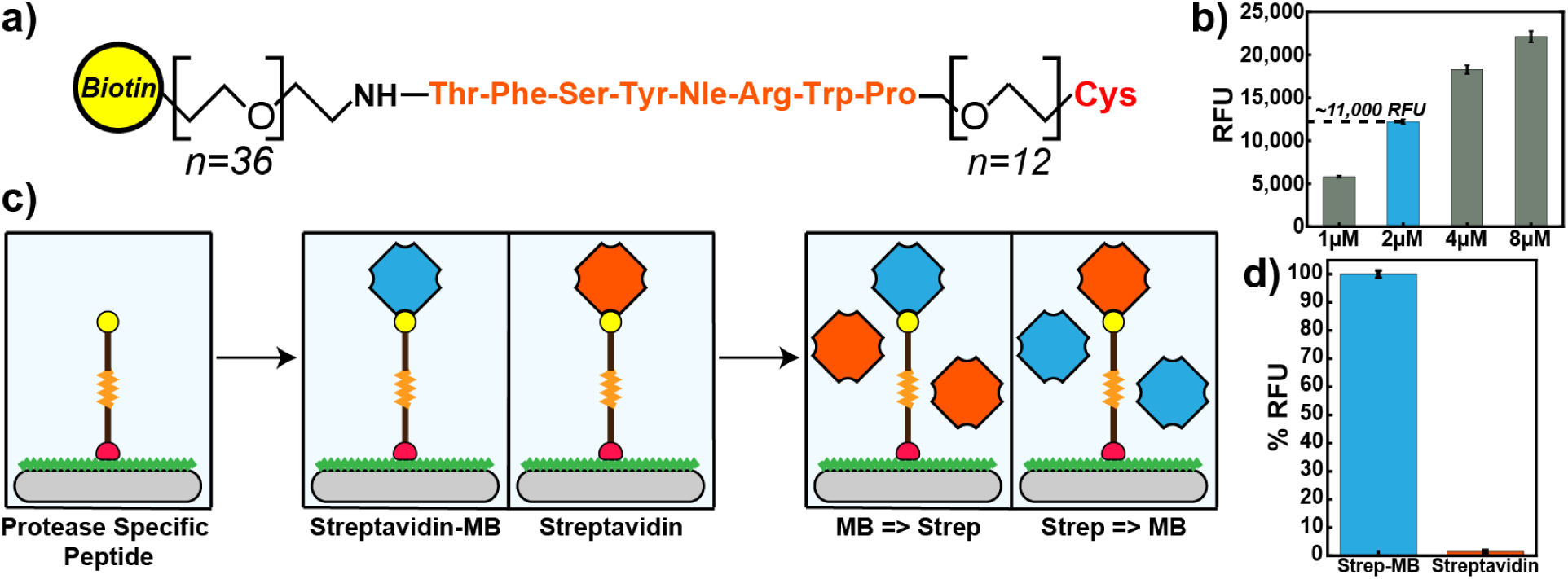
(a) Design of a peptide substrate for a papain heterogeneous assay. Amino acids are listed using the standard three-letter code. (b) Standard curves for peptide loading capacity. A concentration of 2 µM was used for subsequent optical assays yielding an RFU reading of ∼11,000. (c) Illustration of peptide immobilization scheme and streptavidin loading/blocking on polystyrene plates. (d) Bar graphs comparing the loading of fluorescent streptavidin Marina Blue (blue) and blocking with non-fluorescent streptavidin (orange) to immobilized peptide substrate. Error bars are ±1σ.

### Immobilization of papain peptide substrate

The papain substrate (1 - 8 µM) was incubated for 2 hours in a microplate that was functionalized with BSA-maleimide. Unbound peptide was removed by washing and unreacted maleimide was blocked by the addition of free cysteine. The immobilized peptide containing an *N*-terminal biotin group was then quantified by labeling with streptavidin-marina blue (SA-MB). We showed that the concentration of peptide directly correlated with the fluorescent signal (Fig. 2b) and we chose a peptide concentration of 2 µM for the remainder of our studies.

To ensure that labelling of the immobilized peptide occurs through direct interaction between biotin and streptavidin, a competition assay was performed using SA-MB and non-fluorescent streptavidin (Fig. 2c). When the immobilized papain substrate was first incubated with SA-MB and then with streptavidin, a strong fluorescent signal was detected (Fig. 2d). However, when the peptide substrate was first incubated with non-fluorescent streptavidin followed by SA-MB, no fluorescent signal was detected. These data show that the fluorescent signal generated using this heterogeneous assay is due to direct interaction between SA-MB and the immobilized biotin. In addition, these studies show that binding of the SA-MB complex is stable, as the fluorescent signal cannot be reduced even after 1-hour incubation with non-fluorescent streptavidin.

### Papain cleavage of immobilized peptide

After successful peptide loading, cleavage of this sequence by papain and release of SA-MB was evaluated. We incubated the immobilized peptide-SA-MB complex with 20 nM of papain. At defined time intervals between 2.5 and 30 minutes, the enzyme activity was terminated by adding E-64 (Fig. 3). When all time points were acquired, the wells on the microplate were washed to remove peptide-SA-MB that had been enzymatically released from the surface. Any peptide-SA-MB that was still tethered to the surface was then quantified on the fluorescent microplate reader. Enzymatic activity by papain resulted a time-dependent decrease in fluorescence. After only 5 minutes, the fluorescent signal was reduced by ∼50% and an additional 30% of peptide-SA-MB was removed over the next 15 minutes. The reaction was monitored for an additional 10 minutes (30 minutes total), however there was no further change in fluorescence during this time.

**Figure 3.**
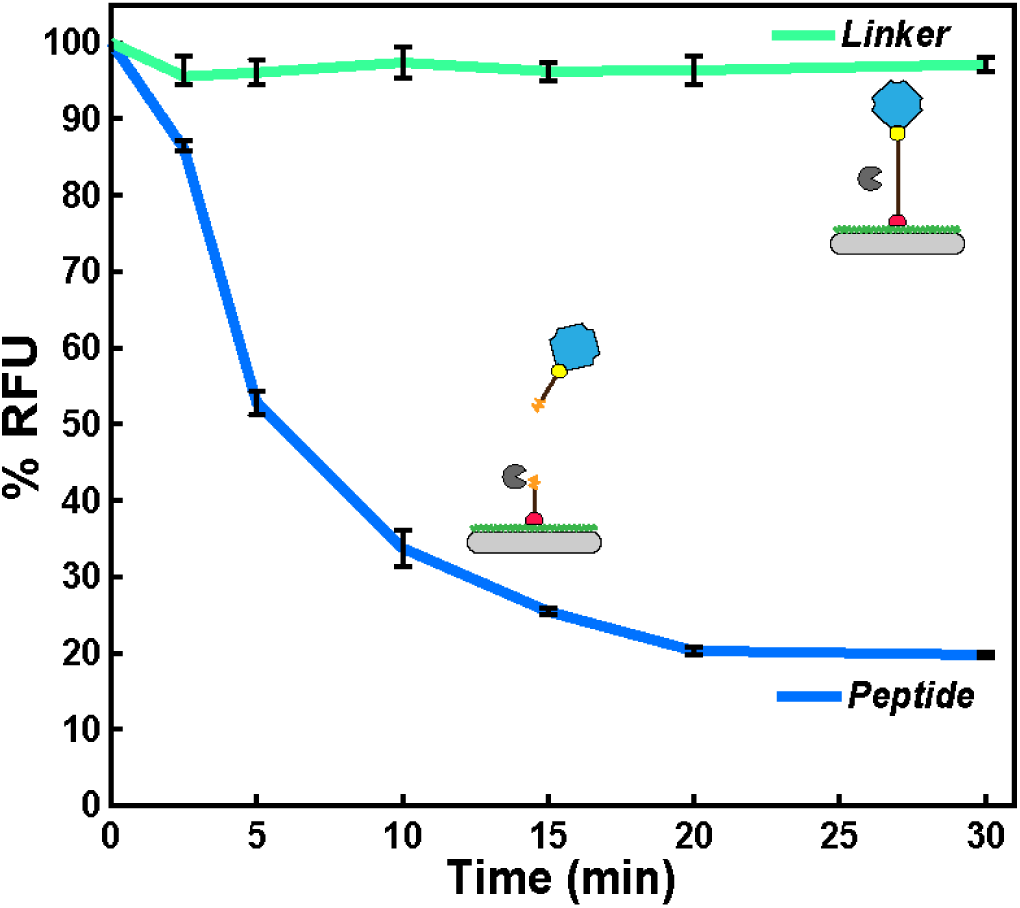
Time-dependent hydrolysis of immobilized peptide substrate by papain. No cleavage of PEG linker was detected under the same conditions. Error bars are ±1σ.

To ensure that the time-dependent reduction in fluorescence was due to the enzymatic activity of papain, a biotin-PEG_75_-thiol (hereafter known as “linker”) was included as a control. This linker lacks peptide bonds and therefore cannot be cleaved by papain; however, it can be immobilized to the plate surface via a covalent thioether linkage and bind to SA-MB. We show that papain was unable to reduce the fluorescence signal in wells that contained this linker, therefore confirming that papain cleavage of the peptide causes the release of SA-MB in this heterogeneous assay.

### Immobilization of peptide substrate to GMR SV sensor

After verifying cleavage of the immobilized peptide by papain, the assay was translated to the GMR SV sensor array to demonstrate its potential for use in a wash-free, real-time, heterogeneous assay. The biochips consist of an 8×10 array of GMR SV sensors each measuring 90 µm^2^ and are coated with a 50 nm SiO_2_ passivation layer to protect them from corrosion^41^. The peptide was first conjugated to BSA-maleimide and this complex was nanospotted onto 25 sensors on the biochip. The biochip was placed into a custom designed measurement station that can detect changes in MR (Fig. S1). Streptavidin-coated MNP (SA-MNP) containing multiple Fe_2_O_3_ cores embedded in a dextran matrix were added. At this size scale, the MNPs are superparamagnetic, and we predicted that when orientated in close proximity to the sensor surface (<200 nm) an increase in MR will occur from the stray field generated by the MNPs.

SA-MNPs were added to the biochip and binding to the biotinylated peptide was monitored in real-time for 32.5 minutes. The change in MR on the peptide spotted sensors (n=16) was rapid in the initial 5-min interval (153.6 ppm/min) and then slowed over the remaining 25 min, from 31.0 ppm/min (between 5-10 min) to 1.34 ppm/min (from 25 – 30 min) (Fig. 4a). Streptavidin was pre-incubated with 7 sensors on the biochip, prior to addition of SA-MNPs. Under these conditions, no change in MR occurred. These data show that the change in MR is due to direct binding of SA-MNPs to the immobilized peptide molecule. This assay format can clearly distinguish between bound and unbound SA-MNPs and therefore will enable us to perform protease-mediated MNP release assays.

**Figure 4.**
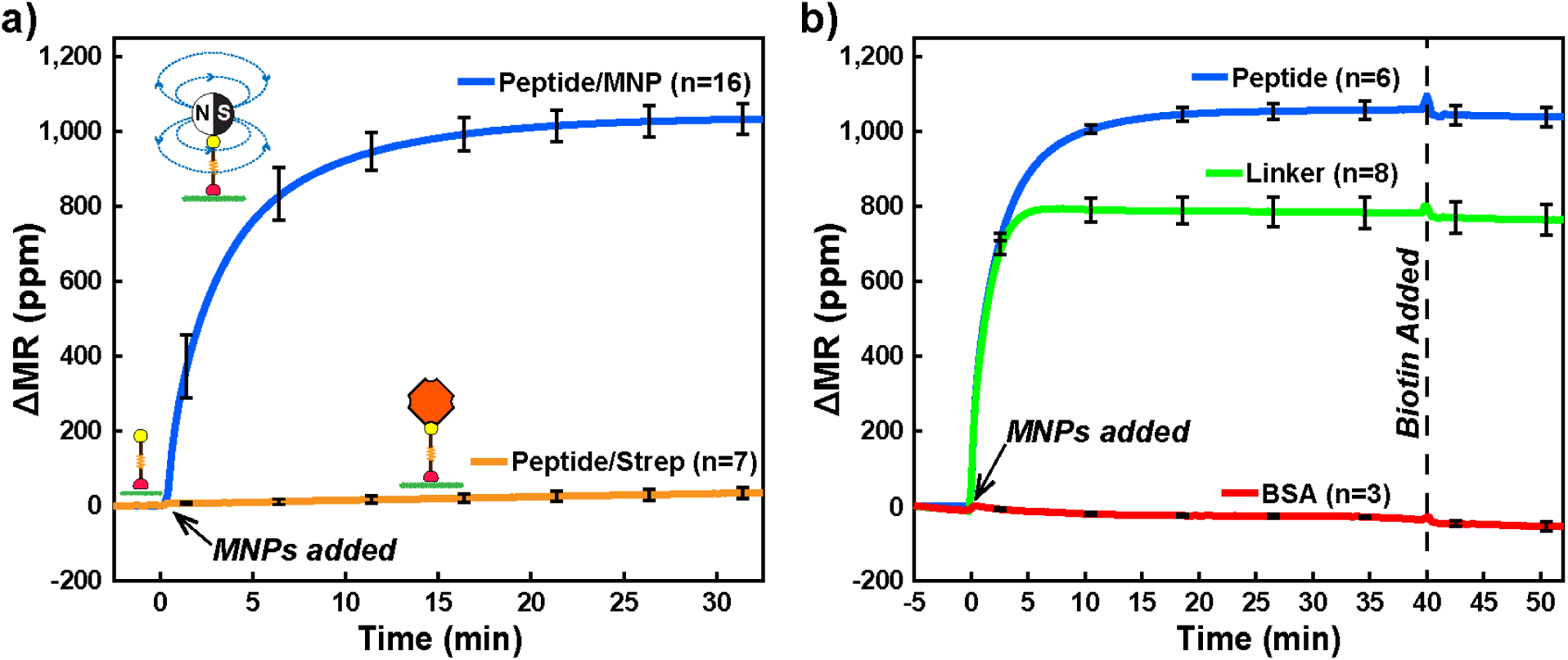
Real-time magnetometry loading data. (a) Loading of MNPs to immobilized peptides that were blocked with streptavidin (orange) or had no blocking (blue). (b) MNP loading to sensors via immobilized peptide (blue) and linker (green). Sensors lacking peptide and linker contained only BSA (red). Curves depicted are the mean signal of sensors and n corresponds to number of sensors that were functionalized. After 40 minutes, all sensors were incubated with biotin to block all available streptavidin sites on MNPs. Error bars are ±1σ and ΔMR is the change in magnetoresistance from the initial value (MR_0_) in parts-per-million (ppm).

Additional control conditions were also evaluated on the GMR SV sensors. We showed that the thiol-PEG_75_-biotin linker can be used to immobilize SA-MNPs to the biochip thereby providing us with a non-cleavable control for papain activity assays (Fig. 4b). After 40 minutes of SA-MNPs loading, the GMR sensor were incubated with excess biotin to block all available streptavidin molecules on the MNP surface. No MNPs were released from the surface indicating that there was strong affinity between the immobilized biotin and streptavidin on the MNPs (Fig. 4b). This blocking step was predicted to be important for downstream papain digestion assays, as SA-MNPs released by the action of the protease would be unable to re-attach to the surface.

### GMR SV-based protease assays

A sensor array containing immobilized SA-MNPs was incubated with 20 nM of papain for 160 minutes and the MNPs were released from the surface in a time-dependent manner. After 160 minutes, 45% of the signal was reduced without any washing-steps being performed at pH 7.4 (Fig. 5a). Under the same conditions, sensors containing the non-cleavable linker sequence, showed only 6% reduction in MNP signal. These studies confirmed that protease activity can be measured in real-time using a wash-free GMR SV sensor assay.

**Figure 5.**
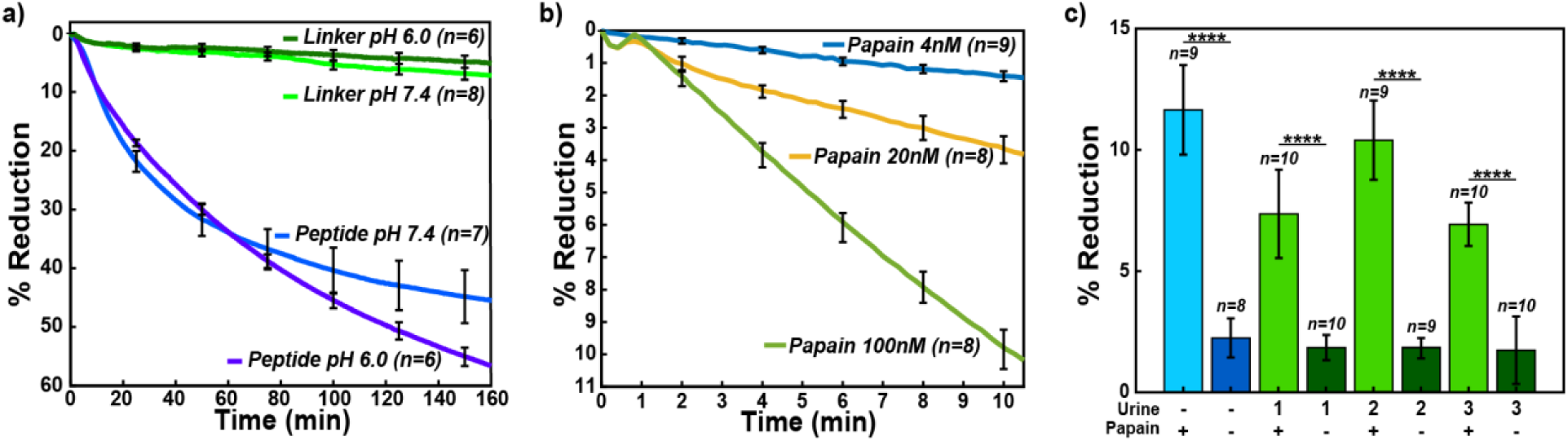
Measured real-time magnetometry papain digestion. (a) Normalized % reduction of peptide and linker sensors treated with 20 nM papain in pH 6.0 and pH 7.4 conditions. (b) Normalized % reduction data comparing the total % reduction for 4 nM (blue), 20 nM (yellow), and 100 nM (green) at pH 7.4 peptide treated sensors. Curves depicted are the mean signal of sensors that were functionalized. (c) Normalized % reduction of papain spiked urine samples and PBS. Error bars are ±1σ.

For a magnetic nanosensor protease assay to have utility in a POC or POU setting, it should be able to rapidly quantify the protease concentration in a biofluid sample under a variety of assay conditions. Biofluids such as plasma, sputum, and wound fluid have a pH value close to neutral, and therefore the prototype assay described above is suitable for detecting protease activity under these conditions. However, other biofluids such as urine have a pH of 6.0^46^, and it was unclear if the assay was compatible with these mildly acidic conditions. Papain is enzymatically active at pH 6.0 and therefore we evaluated the magnetic nanosensor assay in these conditions. We show that 57% of the MNPs are released from the sensor surface after 160 minutes incubation with 20 nM of papain at pH 6.0 while only 4% of the linker is released (Fig. 5a). Papain is more stable at acidic pH than at neutral pH and therefore the greater release of MNPs at pH 6.0 relative to pH 7.4 is likely due to increased stability of the enzyme in the acidic environment.

Next, we determined if the rate of MNP release correlates with enzyme concentration and if sufficient enzyme activity data can be generated within a shorter assay time. To do this, we assayed MNP coated biochips with 4, 20, and 100 nM of papain for 10 minutes. The MNP release from the surface strongly correlates with the concentration of enzyme in the assay (Fig. 5b). Therefore, the rate of change in MR can be used to quantify the amount of papain in the assay. The minimum assay time where protease cleavage can be quantified corresponds to the time when the signal-to-noise ratio (SNR) is greater than 2. Under these conditions, 4 nM of papain was detected after 3.49 minutes incubation while 100 nM of papain can be quantified after 2.87 minutes incubation. These data validate the use of a GMR sensors to rapidly quantify protease activity using a peptide release assay.

### Papain activity in urine

Urine is a commonly used biofluid for diagnosis of urinary tract infections caused by either bacteria or yeast^47^. Proteomic studies have shown that at least 41 proteases are present in urine from healthy individuals and fluorescent reporter peptides have been used to detect activity from these enzymes^48^. The most abundant urine proteases prefer cleaving one or two amino acids from the free N-terminus of proteins and peptides and therefore we predicted that these enzymes would not cleave the Biotin-PEG_36_-TFSYnRWP-PEG_12_-Cys peptide because the N-terminal threonine (T) residue is coupled to PEG_36_ and therefore blocked. Urine also contains a broad acting cysteine protease inhibitor, cystatin C that potently inhibits papain. Using urine from three healthy individuals, cystatin C was inactivated with human cathepsin B. When these urine samples were then added to the GMR-SV sensor containing the peptide and incubated for 10 minutes, no significant reduction of MNPs was detected. This study shows that cathepsin B and the endogenous urine proteases are unable to cleave the papain peptide substrate. Addition of 20 nM papain to these urine samples for 10 minutes, resulted in a reduction in MR that was comparable to papain assayed in PBS (Fig. 5c). The average time for the SNR to be greater than 2 in these samples was 3.76±0.25 minutes.

## DISCUSSION

The goal of this research is to demonstrate that GMR SV sensor-based detection can provide a POC or POU platform to quantify proteases in patient samples. We show that the rate of MNP release by papain directly correlates with the concentration of papain in the assay. Therefore, this platform technology can utilize protease activity in a biological sample as a proxy for quantifying the concentration of the enzyme. Importantly, papain does not release MNPs when these particles are tethered to the sensor surface by a non-peptide linker. In addition, we show that a protease-containing sample such as urine, does not release MNPs when the peptide substrate was designed for cleavage by papain. However, upon addition of papain to the urine, the MNPs are released at a similar rate to reactions that contain just papain in assay buffer.

In the described assay set-up, any peptide substrate that is cleaved by a protease-of-interest can be used in this magnetic sensor assay. The current GMR SV chips contain 80 sensors allowing for quantitation of multiple proteases in the same biofluid sample if they cleave different peptide substrates. These studies demonstrate that bacterial or fungal proteases present in urine of patients with urinary tract infections can be quantified by developing a GMR SV sensor assay containing peptide substrates that are specifically cleaved by the microbial proteases. Using a papain concentration of 4 nM, we were able to detect protease activity in 3.5 min and therefore this assay has the potential to rapidly diagnose urinary tract infections, an important parameter for POC device usage. A magnetic detection strategy leverages the inherent negligible background signal in a biofluid which will allow us to achieve comparable sensitivity in serum, sputum, cyst fluid, semen, and wound fluid as urine.

The novel approach reported here has the potential to address several limitations posed by current protease sensors that have been described. Table 1 shows a comparison between this study and other protease detection strategies based on the buffer and/or biofluid tested, sensitivity, assay time, and sample preparation. Other detection methods such as electrochemical and SERS sensors have design requirements for the peptide substrate such that charged amino acids have to be included for optimal detection^24,27^. Peptides used in SPR sensors must have uncharged amino acids to minimize background signal due to their sensitivity to surface charge^26^. Fluorescent and colorimetric protease assays have been developed to detect protease activity in biofluids such as urine^49,50^, cyst fluid^13^, serum^51,52^, semen^53^, and sputum^54^ as well as used to discover protease inhibitors that have subsequently been developed into drugs^55,56,57,58,59^. However, many of these detection modalities require laboratory equipment, and thus are not suitable for POC and POU applications. GMR SV sensor arrays are compatible with complementary metal-oxide-semiconductor (CMOS) technology which can allow them to be inexpensively mass produced in a disposable format that is amenable to daily use, as well as integration into smartphone-based POC applications^60,61,62^. The sensor can also be produced in a pre-assembled manner by freeze-drying or lyophilizing the complexes such that it is in a “sample-to-answer” format without the need of additional detection reagents used in traditional ELISA assays. Real-time quantification of protease activity using GMR SV sensor-based detection has significant advantages over current POC protease test kits that detect elevated protease activity in wound fluid^63^ or sputum^64^.

**Table 1.**
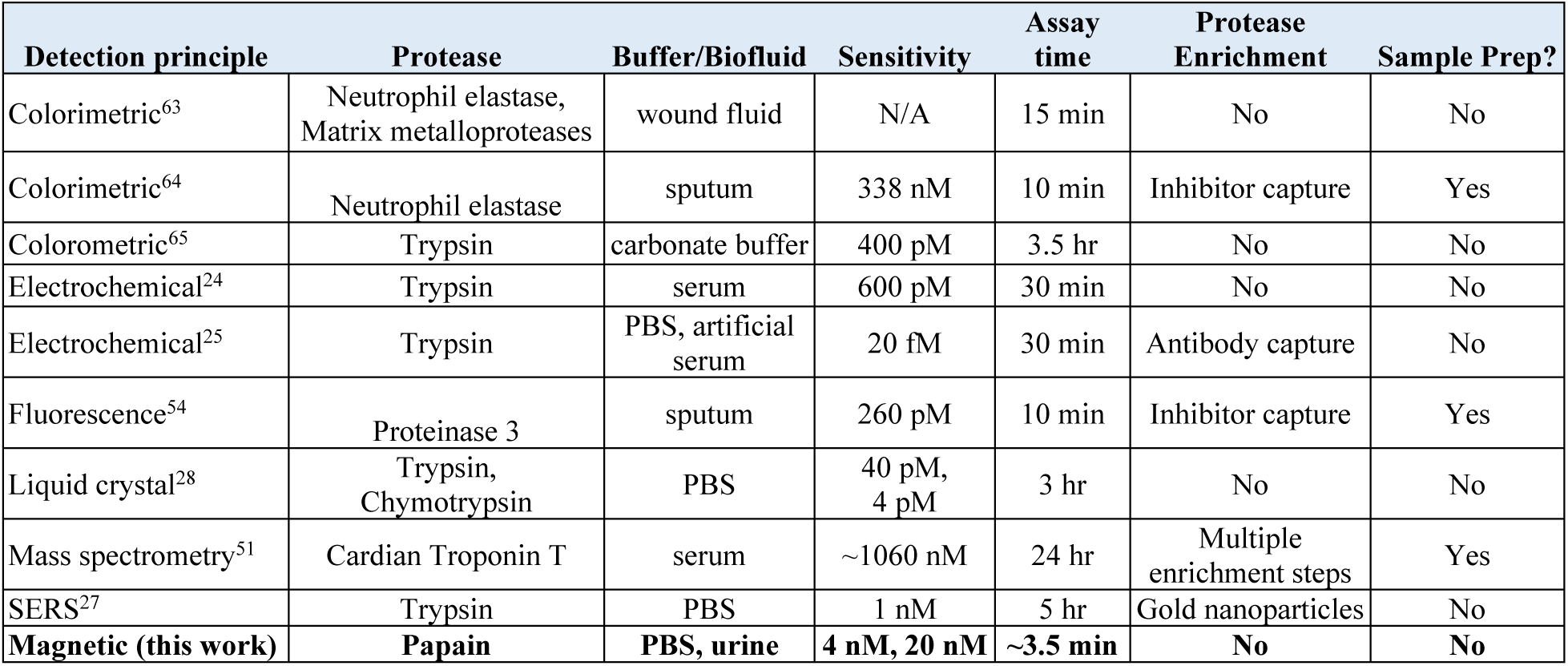
Comparison between several protease-based assays developed and this work.

The long-term goal of this research is to show that GMR SV sensor-based detection can provide a POC platform to detect various disease related proteases in patient samples. Future peptide sensors will explore the use of alternative peptide substrates that are selectively cleaved by either human or microbial protease targets. In addition, if multiple different peptide substrates provide a more accurate read-out for a particular disease, then these substrates can be multiplexed on a single GMR SV chip.

## METHODS

### Peptide/Linker immobilization on optical plates

Biotin-PEG_36_-Thr-Phe-Ser-Tyr-Nle-Arg-Trp-Pro-PEG_12_-Cys (known as peptide) was synthesized by CPC Scientific and Biotin-PEG-SH (known as linker) was purchased from NANOCS. Peptide (2 µM) and linker (128 µM) were prepared in 0.1 M Na_2_HPO_4_, pH 7.4, 0.15 M NaCl, 10 mM EDTA (binding buffer) and added to maleimide activated wells of a 96-well plate (Pierce catalog #15153) and incubated for 2 h. All wells were then blocked with 57 µM Cysteine-HCl (Pierce) in D-PBS for 1 h followed by 1% BSA in D-PBS for 1 h. Wells were washed extensively with 0.1 M Na_2_HPO_4_, pH 7.2, 0.15 M NaCl, 0.05% Tween-20 (buffer A) before and after each step. All steps were performed at room temperature.

### Blocking with streptavidin on optical plates

After peptide/linker immobilization on maleimide coated wells, background fluorescence in each well was measured in a microplate reader with an excitation of 360 nm and an emission of 460 nm. 100 µL of 5 µg/mL Streptavidin, Marina Blue conjugate (ThermoFisher Scientific #S11221) in D-PBS or 20 µg/mL streptavidin (ThermoFisher Scientific #21122) in D-PBS was added to wells and incubated for 1 h. Wells were washed extensively with buffer A and then incubated with 100 µL of 5 µg/mL Streptavidin, Marina Blue conjugate or 20 µg/mL of streptavidin for 1 h. After washing, 100 µL of D-PBS was added to each well and fluorescence was measured. All steps were performed at room temperature. The background fluorescence quantified in each well prior to addition of streptavidin was subtracted from the total fluorescence after addition of streptavidin.

### Time-dependent digestion on optical plates

20 nM of papain in D-PBS, 2 mM DTT was added to each well and the reaction was incubated at room temperature. After defined time intervals of 0, 2.5, 5, 10, 15, 20, and 30 mins, the reaction was stopped by addition of E-64 to a final concentration of 20 µM. Wells were washed extensively with buffer A, hydrated with 100 µL of D-PBS, and fluorescence was measured as outlined above.

### Percent reduction calculations for optical assays

To calculate the reduction of fluorescent signal, the background fluorescence [A] is subtracted from both the pre-assay [B] and post-assay [C] fluorescence; divided by the difference between [A] and [B] and expressed as a percentage.

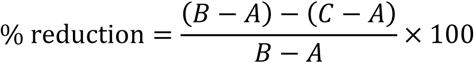

### Conjugation of maleimide activated BSA to peptide and linker

20 µL of 1.33 mg/mL solution of BSA-maleimide (Imject™ Maleimide-Activated BSA, ThermoFisher Scientific #77115) in water, 56 µL of 273 µM peptide or 700 µM linker, and 24 µL of binding buffer were incubated for 24 hours at 4°C then dried in a vacuum centrifuge. This complex was then dissolved in 10 µL of water.

### Chip functionalization

The chip was placed in a UV ozone cleaner for 10 minutes. A custom 3-D printed structure (printed using a Zortrax) with an attached silicone O-ring (High-Temperature High-Purity Silicone O-Rings for Tube Fittings with Straight-Thread Connection, McMaster-Carr #9396K422) was then screwed onto the chip to allow reagents to be pipetted onto the surface without leakage (Fig. S1a). The chip was washed with 100 µL water and 1 µL of BSA (5%), BSA-peptide (1.33 mg/mL), 1.33 mg/mL BSA-Linker) was nanospotted onto the corners of the chip sensors with some separation. The chip was then incubated at 4°C in a humidity chamber for 24 h, washed with water, and then blocked with 100 µL 5% BSA for 30 minutes.

### Magnetic chip reader

The measurement setup consists of a computer, a power amplifier (PA, Kepco 20-5D), a Helmholtz coil and custom readout electronics^66^. The GMR SV sensor chip, which has 80 individually addressable sensors with a nominal resistance of 1.3 kΩ and an MR ratio of 10.6%, was placed firmly inside of the Helmholtz coil using a custom designed sensor holder (Fig. S1b). A double modulation readout scheme was used to reject 1/*f* noise from both the sensors and electronics, and temperature compensation technique was used to reduce temperature drift^67^. The computer digitally adjusted the frequencies and amplitudes of sensor bias voltage and magnetic field through a National Instruments data acquisition card (DAQ, NI PCIe-6351) and a LabVIEW graphical user interface (GUI). Specifically, the PA controlled by the computer provided current into the Helmholtz coil, which offers homogenous magnetic field (23 – 34 Oe_rms_ based on the sensor MR) for the sensor chip. The readout electronics contain 8× transimpedance amplifiers to convert the currents to voltages that was quantized by the DAQ. Time-multiplexing was applied to read out the entire 8×10 sensor array with a 10 second update rate.

### Magnetic sensor assays

1 µL of 1 mg/mL streptavidin was nanospotted onto sensors containing BSA, BSA-peptide, or BSA-linker and incubated for 30 minutes followed by washing with water and then PBS. The chip was then connected to the electromagnet measurement station (Fig. S1B). Sensors were loaded with 60 µL of 50 nm magnetic nanoparticles coated with streptavidin (Miltenyi Biotec #130-048-101) for 35 minutes followed by five washes with 100 µL buffer A. 100 µL of 1 mM Biotin (Fisher Scientific, 15486) was added for 10 minutes followed by five washes with 100 µL buffer A. 100 µL of papain in D-PBS, pH 7.4 or D-PBS, pH 6.0 (adjusted to pH 6.0 with citric acid) was added and incubated for up to 200 minutes. To stop the assay, the sensor was washed five times in buffer A.

### Papain activity in urine

Urine from health donors was purchased from Innovative Research, Novi, Michigan (IRHUURES50ML). 97 µL of urine was incubated with 1 µL of human cathepsin B (R&D Systems, 953-CY-010) and 1 µL of DTT for 15 minutes to inactivate the urine cysteine protease inhibitor, cystatin C. The cystatin C-inactivated urine was then added to the sensor well and magnetoresistance was evaluated for 20 minutes. 1 µL of papain was then added to the urine and magnetoresistance was evaluated for 100 minutes. The final concentration of cathepsin B, DTT, and papain in the urine assay was 10 µM, 2 mM, and 10 nM, respectively.

### Magnetic assay time of detection of papain activity calculations

The signal-to-noise ratio (SNR) for each time point was determined by calculating the difference of mean signal (S) for the peptide and linker sensors divided by the noise which is equal to square root of the sum of variances (σ^2^) of peptide and linker data.

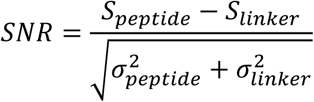

## Acknowledgements

X.Z. and D.A.H acknowledge funding from the National Science Foundation (Grant ECCS-1454608). The research was supported by the National Institutes of Health, Grant UL1TR001442 and EB028485 awarded to A.J.O and D.A.H.

## Author contributions statement

S.A., S.J., A.J.O., and D.A.H. conceived the research. S.A., S.J., and M.S. designed and conducted the optical and magnetic assay experiments. X.Z. designed the electronic system and signal processing. S.A., S.J., M.S, X.Z., A.J.O., and D.A.H. wrote the paper. All authors discussed the results and reviewed the manuscript.

## Additional information

### Competing financial interests

D.A.H. has related patents or patent applications assigned to Stanford University and out-licensed for potential commercialization. D.A.H. has stock in MagArray, Inc. and Flux Biosciences, which have licensed relevant patents from Stanford University for commercialization of GMR biosensor chips.

---

**Figure S1.**
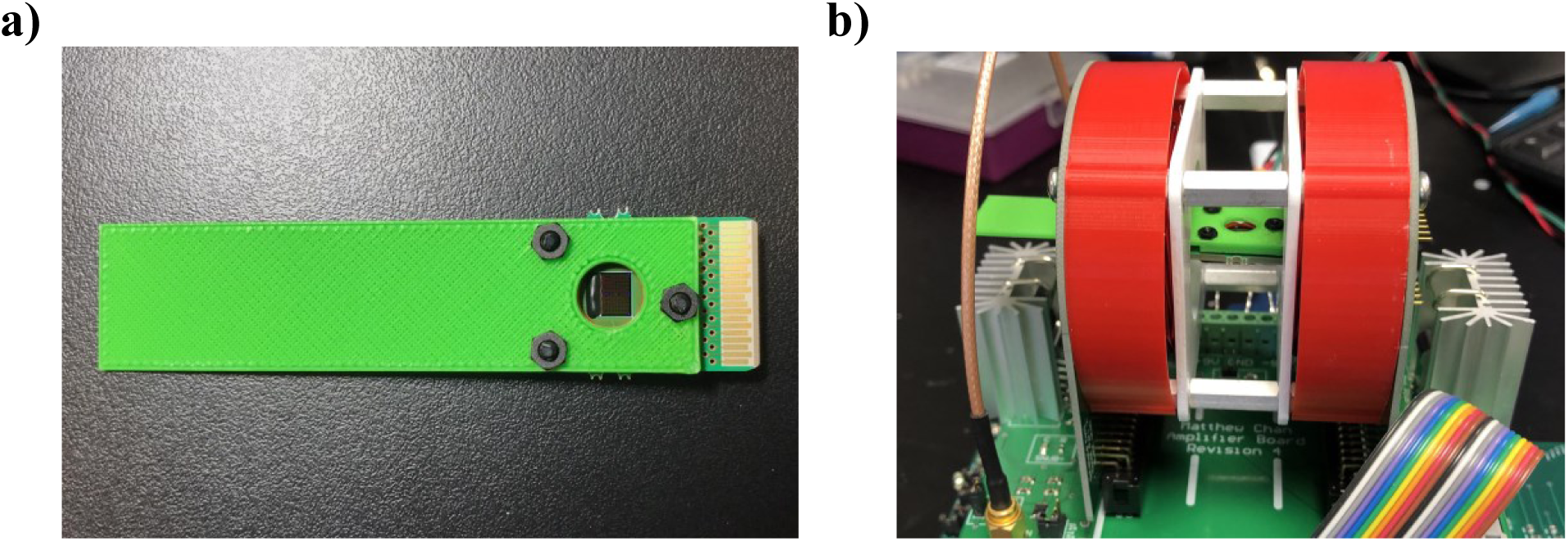
Magnetic assay setup. a) 3-D printed holder screwed to chip creating a 100 µL reservoir over biochip sensor array to allow reagents to be pipetted onto the surface. b) Chip plugged into electromagnet to begin real-time magnetometry measurements.

## REFERENCES

1. Whitcomb, D. C. & Lowe, M. E. Human pancreatic digestive enzymes. Dig. Dis. Sci. 52, 1–17 (2007).

2. Janis, J. E. & Attinger, C. E. The basic science of wound healing. Plast. Reconstr. Surg. 117, 12S–34S (2006).

3. Heutinck, K. M., ten Berge, I. J., Hack, C. E., Hamann, J. & Rowshani, A. T. Serine proteases of the human immune system in health and disease. Mol. Immunol. 47, 1943–1955 (2010).

4. Hochstrasser, M. Ubiquitin, proteasomes, and the regulation of intracellular protein degradation. Curr. Opin. Cell Biol. 7, 215–223 (1995).

5. Duan, G. & Walther, D. The Roles of Post-translational Modifications in the Context of Protein Interaction Networks. PLoS Comput. Biol. 11, (2015).

6. López-Otín, C. & Bond, J. S. Proteases: multifunctional enzymes in life and disease. J. Biol. Chem. (2008).

7. Martinelli, P. & Rugarli, E. I. Emerging roles of mitochondrial proteases in neurodegeneration. Biochim. Biophys. Acta 1797, 1–10 (2010).

8. Trindade, F., Ferreira, R., Amado, F. & Vitorino, R. Biofluid proteases profiling in diabetes mellitus. Adv. Clin. Chem. 69, 161–207 (2015).

9. Musante, L. et al. Proteases and protease inhibitors of urinary extracellular vesicles in diabetic nephropathy. J. Diabetes Res. 2015, 289734 (2015).

10. Stamey, T. A. et al. Prostate-Specific Antigen as a Serum Marker for Adenocarcinoma of the Prostate. N. Engl. J. Med. 317, 909–916 (1987).

11. Sabbagh, B. et al. Functional protease profiling for laboratory based diagnosis of invasive aspergillosis. Int. J. Oncol. 47, 143–150 (2015).

12. Findeisen, P. & Neumaier, M. Functional protease profiling for diagnosis of malignant disease. Proteomics Clin. Appl. 6, 60–78 (2012).

13. Ivry, S. L. et al. Global Protease Activity Profiling Provides Differential Diagnosis of Pancreatic Cysts. Clin. Cancer Res. 23, 4865–4874 (2017).

14. MacNee, W. Pathogenesis of Chronic Obstructive Pulmonary Disease. Proc. Am. Thorac. Soc. 2, 258–266 (2005).

15. Barnes, P. J., Shapiro, S. D. & Pauwels, R. A. Chronic obstructive pulmonary disease: molecular and cellularmechanisms. Eur. Respir. J. 22, 672–688 (2003).

16. Voynow, J. A., Fischer, B. M. & Zheng, S. Proteases and cystic fibrosis. Int. J. Biochem. Cell Biol. 40, 1238–1245 (2008).

17. Birrer, P. et al. Protease-antiprotease imbalance in the lungs of children with cystic fibrosis. Am. J. Respir. Crit. Care Med. 150, 207–213 (1994).

18. Quinn, R. A. et al. Neutrophilic proteolysis in the cystic fibrosis lung correlates with a pathogenic microbiome. Microbiome 7, 23 (2019).

19. Guo, S. & DiPietro, L. A. Factors Affecting Wound Healing. J. Dent. Res. 89, 219–229 (2010).

20. McCarty, S. M. & Percival, S. L. Proteases and Delayed Wound Healing. Adv. Wound Care 2, 438–447 (2013).

21. Ong, I. L. H. & Yang, K.-L. Recent developments in protease activity assays and sensors. The Analyst 142, 1867–1881 (2017).

22. Parks, T. D., Leuther, K. K., Howard, E. D., Johnston, S. A. & Dougherty, W. G. Release of proteins and peptides from fusion proteins using a recombinant plant virus proteinase. Anal. Biochem. 216, 413–417 (1994).

23. Callahan, B. P., Stanger, M. J. & Belfort, M. Protease activation of split green fluorescent protein. Chembiochem Eur. J. Chem. Biol. 11, 2259–2263 (2010).

24. Cao, Y., Yu, J., Bo, B., Shu, Y. & Li, G. A simple and general approach to assay protease activity with electrochemical technique. Biosens. Bioelectron. 45, 1–5 (2013).

25. Park, S., Kim, G., Seo, J. & Yang, H. Ultrasensitive Protease Sensors Using Selective Affinity Binding, Selective Proteolytic Reaction, and Proximity-Dependent Electrochemical Reaction. Anal. Chem. 88, 11995–12000 (2016).

26. Chen, H. et al. Potassium ion sensing using a self-assembled calix[4]crown monolayer by surface plasmon resonance. Sens. Actuators B Chem. 133, 577–581 (2008).

27. Yang, L. et al. SERS determination of protease through a particle-on-a-film configuration constructed by electrostatic assembly in an enzymatic hydrolysis reaction. RSC Adv. 6, 90120–90125 (2016).

28. Chen, C.-H. & Yang, K.-L. Oligopeptide immobilization strategy for improving stability and sensitivity of liquid-crystal protease assays. Sens. Actuators B Chem. 204, 734–740 (2014).

29. McFadyen, I. R., Fullerton, E. E. & Carey, M. J. State-of-the-Art Magnetic Hard Disk Drives. MRS Bull. 31, 379–383 (2006).

30. Chappert, C., Fert, A. & Van Dau, F. N. The emergence of spin electronics in data storage. in Nanoscience and Technology 147–157 (Co-Published with Macmillan Publishers Ltd, UK, 2009). doi: 10.1142/9789814287005_0015.

31. Spong, J. K., Speriosu, Fontana R. E., Dovek, M. M. & Hylton, T. L. Giant magnetoresistive spin valve bridge sensor. IEEE Trans. Magn. 32, 366–371 (1996).

32. Pelegri, J. et al. A novel spin valve bridge sensor for current sensing. in IMTC 2001. Proceedings of the 18th IEEE Instrumentation and Measurement Technology Conference. Rediscovering Measurement in the Age of Informatics (Cat. No.01CH 37188) vol. 1 422–424 vol.1 (2001).

33. Daughton, J. M. Magnetic tunneling applied to memory (invited). J. Appl. Phys. 81, 3758–3763 (1997).

34. Baselt, D. R. et al. A biosensor based on magnetoresistance technology1This paper was awarded the Biosensors & Bioelectronics Award for the most original contribution to the Congress.1. Biosens. Bioelectron. 13, 731–739 (1998).

35. Graham, D. L., Ferreira, H. A. & Freitas, P. P. Magnetoresistive-based biosensors and biochips. Trends Biotechnol. 22, 455–462 (2004).

36. Parkin, S. S. P. Origin of enhanced magnetoresistance of magnetic multilayers: Spin-dependent scattering from magnetic interface states. Phys. Rev. Lett. 71, 1641–1644 (1993).

37. Aubin, J. E. Autofluorescence of viable cultured mammalian cells. J. Histochem. Cytochem. Off. J. Histochem. Soc. 27, 36–43 (1979).

38. Giloh, H. & Sedat, J. W. Fluorescence Microscopy: Reduced Photobleaching of Rhodamine and Fluorescein Protein Conjugates by n-Propyl Gallate. Science 217, 1252–1255 (1982).

39. Lee, J.-R. et al. Multiplex giant magnetoresistive biosensor microarrays identify interferon-associated autoantibodies in systemic lupus erythematosus. Sci. Rep. 6, 27623 (2016).

40. Gaster, R. S. et al. Matrix-insensitive protein assays push the limits of biosensors in medicine. Nat. Med. 15, 1327–1332 (2009).

41. Osterfeld, S. J. et al. Multiplex protein assays based on real-time magnetic nanotag sensing. Proc. Natl. Acad. Sci. 105, 20637–20640 (2008).

42. Costa, A. G., Cusano, N. E., Silva, B. C., Cremers, S. & Bilezikian, J. P. Cathepsin K: its skeletal actions and role as a therapeutic target in osteoporosis. Nat. Rev. Rheumatol. 7, 447–456 (2011).

43. Choe, Y. et al. Development of α-keto-based inhibitors of cruzain, a cysteine protease implicated in Chagas disease. Bioorg. Med. Chem. 13, 2141–2156 (2005).

44. Kim, M. J. et al. Crystal structure of papain-E64-c complex. Binding diversity of E64-c to papain S2 and S3 subsites. Biochem. J. 287, 797–803 (1992).

45. Lapek, J. D. et al. Quantitative Multiplex Substrate Profiling of Peptidases by Mass Spectrometry. Mol. Cell. Proteomics MCP (2019) doi: 10.1074/mcp.TIR118.001099.

46. Welch, A. A., Mulligan, A., Bingham, S. A. & Khaw, K.-T. Urine pH is an indicator of dietary acid-base load, fruit and vegetables and meat intakes: results from the European Prospective Investigation into Cancer and Nutrition (EPIC)-Norfolk population study. Br. J. Nutr. 99, 1335–1343 (2008).

47. Najeeb, S. et al. Comparison of Urine Dipstick Test with Conventional Urine Culture in Diagnosis of Urinary Tract Infection. 25, 3 (2015).

48. Taylor, J. M. et al. Aminopeptidase Activities as Prospective Urinary Biomarkers for Bladder Cancer. Proteomics Clin. Appl. 8, 317–326 (2014).

49. Poon, C.-Y. et al. FRET-based modified graphene quantum dots for direct trypsin quantification in urine. Anal. Chim. Acta 917, 64–70 (2016).

50. Zhao, L. et al. Fluorescent Strips of Electrospun Fibers for Ratiometric Sensing of Serum Heparin and Urine Trypsin. ACS Appl. Mater. Interfaces 9, 3400–3410 (2017).

51. Streng, A. S. et al. Development of a targeted selected ion monitoring assay for the elucidation of protease induced structural changes in cardiac troponin T. J. Proteomics 136, 123–132 (2016).

52. Lin, T. et al. A sensitive colorimetric assay for cholesterol based on the peroxidase-like activity of MoS2 nanosheets. Microchim. Acta 184, 1233–1237 (2017).

53. Etheridge, T., Straus, J., Ritter, M. A., Jarrard, D. F. & Huang, W. Semen AMACR protein as a novel method for detecting prostate cancer. Urol. Oncol. Semin. Orig. Investig. 36, 532.e1-532.e7 (2018).

54. Ferguson, T. E. G. et al. P111 Quantification of active proteinase 3 in sputum samples using a novel activity-based immunoassay. Thorax 73, A162–A162 (2018).

55. Matayoshi, E. D., Wang, G. T., Krafft, G. A. & Erickson, J. Novel fluorogenic substrates for assaying retroviral proteases by resonance energy transfer. Science 247, 954–958 (1990).

56. Kisselev, A. F. & Goldberg, A. L. Monitoring Activity and Inhibition of 26S Proteasomes with Fluorogenic Peptide Substrates. in Methods in Enzymology vol. 398 364–378 (Academic Press, 2005).

57. Moss, M. L. & Rasmussen, F. H. Fluorescent substrates for the proteinases ADAM17, ADAM10, ADAM8, and ADAM12 useful for high-throughput inhibitor screening. Anal. Biochem. 366, 144–148 (2007).

58. Lai, K. S., Ho, N.-H., Cheng, J. D. & Tung, C.-H. Selective Fluorescence Probes for Dipeptidyl Peptidase Activity - Fibroblast Activation Protein and Dipeptidyl Peptidase IV. Bioconjug. Chem. 18, 1246–1250 (2007).

59. Watzke, A. et al. Selective activity-based probes for cysteine cathepsins. Angew. Chem. Int. Ed Engl. 47, 406–409 (2008).

60. Choi, J. et al. Portable, one-step, and rapid GMR biosensor platform with smartphone interface. Biosens. Bioelectron. 85, 1–7 (2016).

61. Ng, E., Yao, C., Shultz, T. O., Ross-Howe, S. & Wang, S. X. Magneto-nanosensor smartphone platform for the detection of HIV and leukocytosis at point-of-care. Nanomedicine Nanotechnol. Biol. Med. 16, 10–19 (2019).

62. Gaster, R. S., Hall, D. A. & Wang, S. X. nanoLAB: an ultraportable, handheld diagnostic laboratory for global health. Lab. Chip 11, 950–956 (2011).

63. Serena, D. T. et al. Preliminary results: Testing for elevated protease activity in clinical practice. Wound Repair Regen. 20, A112.

64. McCafferty, D. F. et al. P116 Neatstik^®^ – a novel point of care test for the measurement of active neutrophil elastase in patients with respiratory disease. Thorax 72, A146–A146 (2017).

65. Ding, X. & Yang, K.-L. Enzymatic Deposition of Silver Particles for Detecting Protease Activity. Part. Part. Syst. Charact. 31, 1300–1306 (2014).

66. Hall, D. A. et al. GMR biosensor arrays: a system perspective. Biosens. Bioelectron. 25, 2051–2057 (2010).

67. Hall, D. A., Gaster, R. S., Osterfeld, S. J., Murmann, B. & Wang, S. X. GMR Biosensor Arrays: Correction Techniques for Reproducibility and Enhanced Sensitivity. Biosens. Bioelectron. 25, 2177–2181 (2010).

